# Signatures of Motion: Decomposition of Adaptive Morphing Flight in Harris’ Hawks

**DOI:** 10.1101/2025.06.03.657611

**Authors:** Lydia A. France, James Shelton, Marco Klein Heerenbrink, Caroline Brighton, Graham K. Taylor

## Abstract

Birds outperform engineered aircraft with exceptional maneuverability, achieved by continuously morphing their wings and tails in flight. Yet the coordination and control of these shape changes remain poorly understood. Using high-speed motion capture of Harris’ hawks, we analyzed 289,000 wing-tail configurations in over 2000 flights and identified four fundamental shape change patterns, or “morphing shape modes”, that capture over 96% of wing and tail variation. Further modes reflect subtle but critical fine-tuning, in line with known morphing control mechanics. The hawks’ morphing flight is highly structured yet flexible, and we find adaptive strategies in response to obstacles, added weight, with maturity, while each individual shows unique morphing signatures. Our approach defines a shared kinematic morphospace for hawk flight, and more broadly a framework that enables future comparative biomechanics, bio-inspired design, and for interpreting high-dimensional natural motion.

## Introduction

Bird wings and tails fold, flex, and twist in flight, allowing agile and reactive control within complex and unpredictable environments [1, 2, 3]. Despite centuries of fascination, the principles governing coordinated wing-tail shape changes and their use in flight control remain largely unknown [4]. Traditional analyses treat morphing flight as a set of independent angles and span changes, making it challenging to understand the critical couplings at play [5, 6, 7]. Moreover, datasets of morphing across different behaviors from free-flying birds remain rare, with studies commonly using cadavers or focus on a few snapshots of wings in flight [8, 9, 10, 11, 12].

Unlike engineered aircraft, birds do not have discrete control surfaces like rudders or spoilers and the key control mechanics remain unclear [13]. The wings and tail are coordinated together to produce asymmetric and symmetric whole-shape adjustments in flight (Fig. 1) [14, 15, 16]. Full surface reconstructions are challenging to record at high volume, and so researchers commonly predefine control surfaces by measuring key angles from the wings. Such angles are non-independent to calculate and cannot be compared across different morphologies. Finding fine-tuned control and couplings are also complex as angles are highly sensitive to small measurement errors [17].

**Figure 1.**
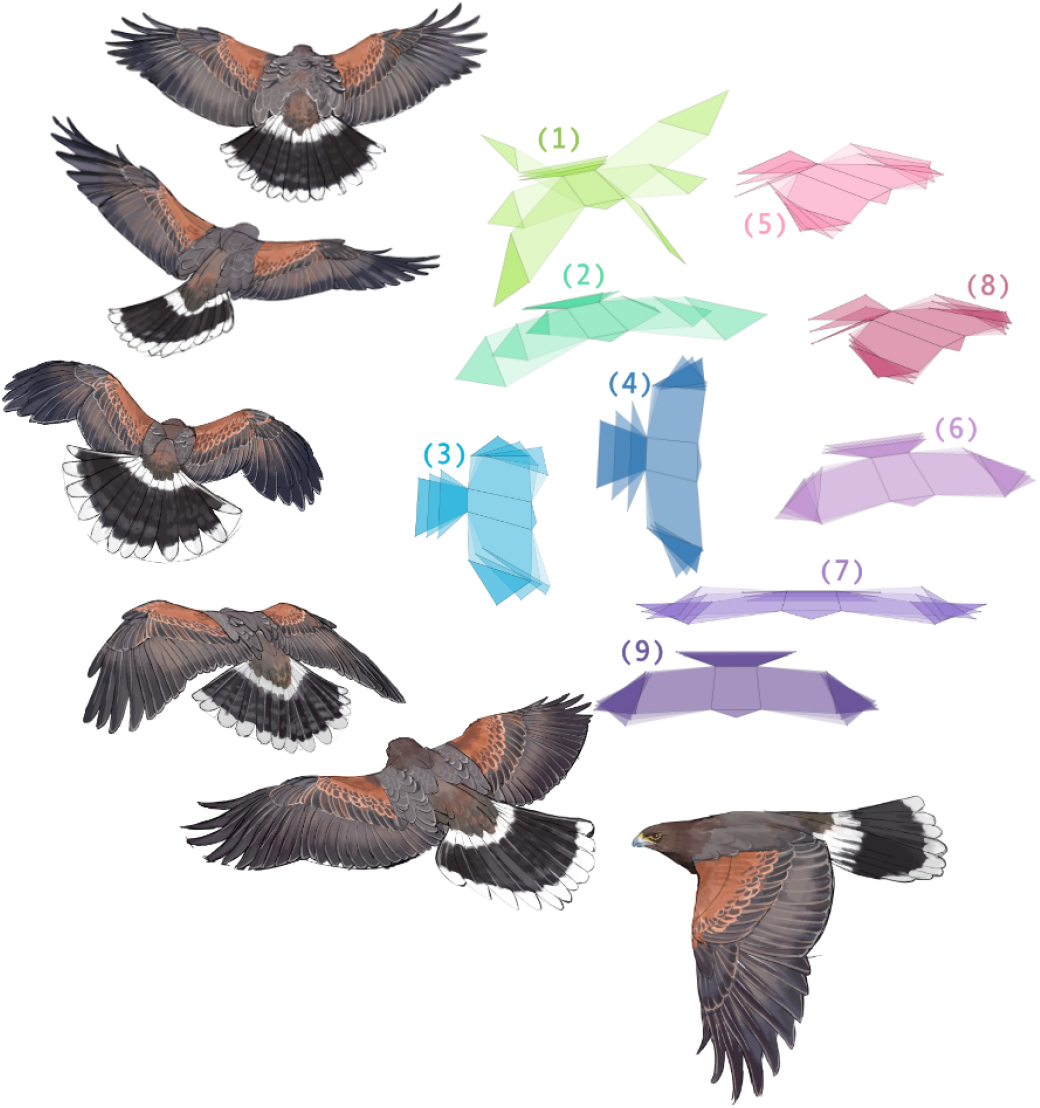
Coordinated wing-tail kinematics split into morphing shape modes. Left: Illustration of a Harris hawk performing a turn and landing. There are no clear control surfaces in bird flight, with complex wing-tail shape changes coordinated with whole body rotations during maneuvers (11). Right: Variation in the hawks’ kinematics can be almost completely represented by combining nine morphing shape modes. Each linear mode, or “eigenbird”, represents highly correlated movements across the wing and tail. Their effect on the mean shape shown in positive and negative directions. Named for their dominant shape change and numbered by variance explained: (1-4) wing lifting, spreading, sweeping and tail spreading; (6,7,9) hand-wing spreading, M-folding, sweeping; (5,8) counter and collective pitching. The effect of each mode is shown here as symmetrical.

We instead use a data-driven approach to study flight kinematics of Harris’ hawks. Rather than a series of executed joint angle changes, we treat morphing flight as highly correlated wing-tail shape changes [4, 18]. We collected an unprecedented dataset of hawk flight with a diverse range of behaviors from five individuals, including flapping, gliding, turning, landing, and obstacle avoidance in a large flight arena. We reconstructed over 280,000 wing-tail shape configurations with motion capture under experimental conditions that varied perch-perch distances, an added obstacle, a worn weight, with experienced and inexperienced fliers [19, 20].

Rather than analyzing isolated movements, we describe a structured kinematic morphospace, a lower-dimensional representation of feasible wing-tail configurations during morphing flight. The kinematic morphospace is defined directly from measured marker positions rotated to the body-axis frame and using principal component analysis [21, 22]. This representation is linear, directly interpretable, and offers a natural language of coordination and flexibility in morphing flight. By projecting the five individuals from different experiments into the same shared space, this allows for direct quantitative comparison which would be otherwise limited in traditional kinematics, revealing the subtle kinematic differences in adaptive flight.

We introduce a low-dimensional decomposition of morphing flight that is robust to noise, stable across individuals, and flexible to different behavioral contexts. This framework offers a tractable and generalizable approach for analyzing and comparing adaptive morphing strategies in avian flight.

## Morphing Shape Change Modes

We identified four dominant morphing shape modes that together explain over 96% of the total variation in wing and tail configuration. When projected onto the mean shape, each mode represents a distinct, coordinated shape change: wing lifting, wing spreading, wing sweeping, and tail spreading (Fig. 1, modes 1-4). Each morphing shape mode defines a set of strongly correlated movements across the wings and tail. For instance, wing lifting (mode 1) co-occurs with tail lifting, a pattern consistent with passive body oscillation. Wing sweeping and tail spreading both co-occur in modes 3 and 4, patterns not mechanically imposed but reliably emerge, consistent with known kinematic patterns previously linked to maneuvering [17]. To test robustness, we applied shuffling, marker sub-sampling, and projection consistency across individuals. The first four PCA modes remained stable in orientation and explained variance even under substantial perturbation. Shuffle controls confirmed that these modes do not emerge from randomly distributed marker coordinates, and reconstruction analysis with four modes showed average marker error within a flight feather width (*<*2 cm).

Further morphing shape modes (modes 5–9) describe subtle adjustments, such as hand-wing sweeping and incidence angle changes in both wings and tail. These smaller variations show greater individual differences and likely contribute to aerodynamic fine-tuning, complementing larger shape changes. Similar patterns such as hand-wing spreading have been observed in flight mechanics of specific maneuvers [23, 2, 7, 24, 25]. Together with the major modes, these account for over 99% of morphing variation by the hawks.

The morphing shape modes emerge as highly correlated spatial relationships between markers on feathers of the hawks in flight (Fig. 2). While birds could use a vast number of configurations by using the full range of all possible joint and muscle positions, in practice maneuvering flight occupies a smaller kinematic subset. After removing whole body rotations and relative to a central body position, we observe wingtip markers formed consistent circular traces during flapping (frontal view) with folding (dorsal view), while tail motion was more limited (lateral view). Even across tens of thousands of examples (see Fig. 2: e.g. 18,556 frames from “Toothless”) marker positions clustered into compact, repeated patterns of shape configurations.

**Figure 2.**
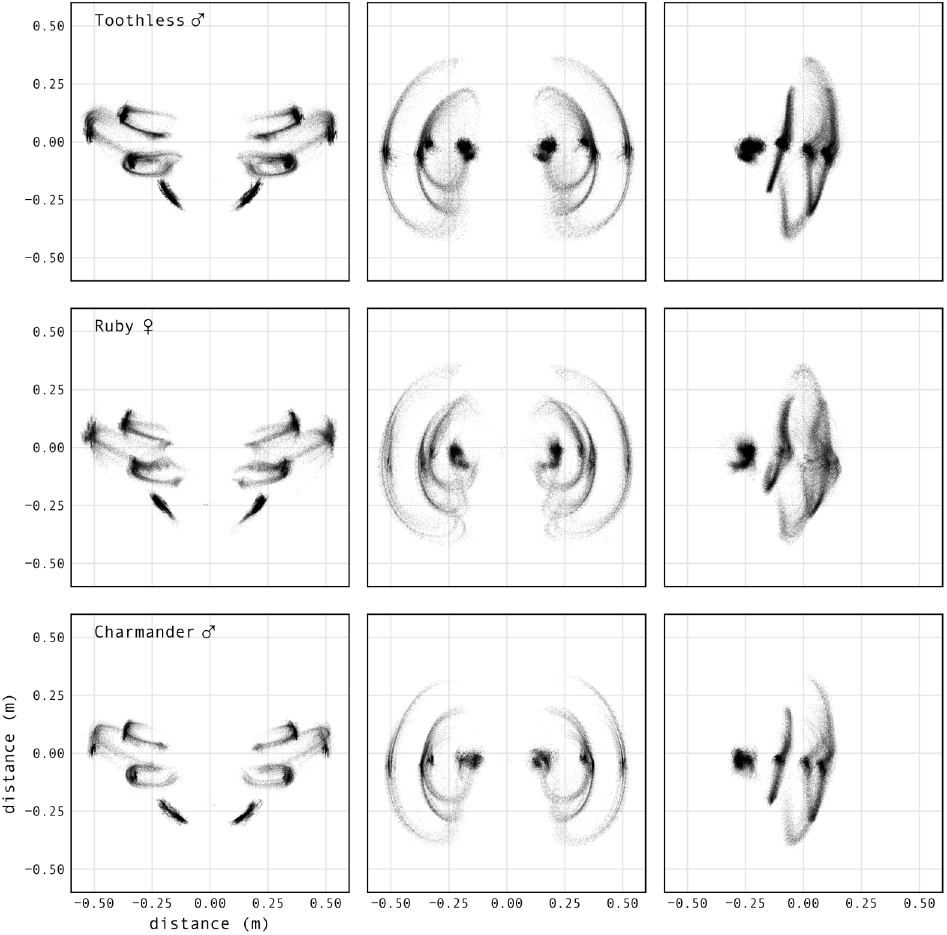
Motion fingerprint in hawk flights. Feather marker positions from three individuals including turning and straight flights, shown from the frontal, dorsal, and lateral views (columns). Dorsal markers on the wings and tail were reconstructed from motion capture and shown in a rotated body-axis system. Showing over a hundred flights overlaid for each hawk, behavior shown here includes flapping, gliding, turning, and landing behaviors. Despite the wings and tail having high degrees of freedom and data including varying flight behaviors, morphing is constrained to a narrow envelope of coordinated shape changes. Data are from 9m flights with and without an obstacle. Rows show: experienced male “Toothless” (18,556 frames, 133 flights); experienced female “Ruby” (12,598 frames, 130 flights); and inexperienced male “Charmander” (9,383 frames, 121 flights).

The hawks’ highly complex morphing flight then collapses to a few linear axes of variation. While PCA is widely used in dimensionality reduction [26], projection back to spatial coordinates provides intuitive decomposition of morphing flight. We can readily compare the differences in morphing in flights with and without an obstacle (Fig. 3). The projections, or scores of the morphing shape modes, each describes coordinated shape change across many markers, with a score of zero the mean shape [22]. As the spatial relationships between the markers are preserved, these results are more intuitive than most examples of PCA. For example, a dihedral pose has a positive wing lifting score, a single variable representing the underlying shape change from multiple markers. We show the binned means and ±1 standard deviation of four individuals with over 250 flights for each experimental condition: with and without an obstacle at the midpoint between two perches. Oscillations are visible during flapping and contrasting with the gliding phase, where wing and tail spreading increases with increased wing pitch on approach to the destination perch. With an added obstacle at the midpoint, the hawks execute a wing sweep maneuver and hand-wing sweep, plus tail banking which is represented by the trough in collective pitching.

**Figure 3.**
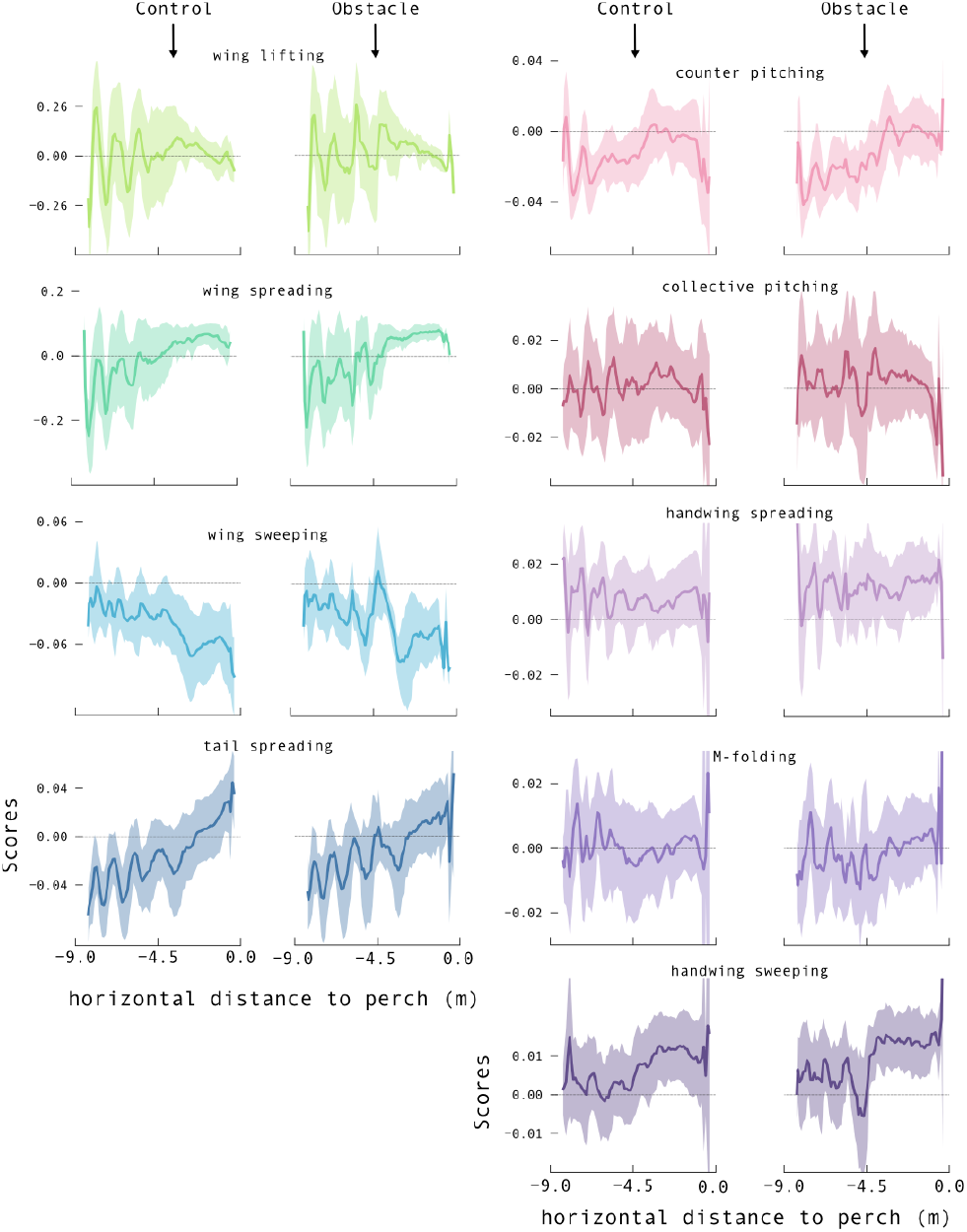
Morphing shape changes with and without an obstacle. Morphing shape changes are shown for 9m flights by four hawks, with and without an obstacle, including flapping, turning, gliding, and landing. Each hawk is projected into a shared kinematic morphospace, allowing direct comparison between individuals and experimental conditions. Differences in flapping and gliding are seen, while subtle kinematics for example wing sweep, tail pitching, and hand-wing adjustments increase near the obstacle. Data represent control flights (N=256; 29,949 frames) and obstacle flights (N=254; 18,426 frames), with the obstacle placed at –4.5m. Morphing shape scores are averaged in 0.1m bins across horizontal distance to perch, with ±1s.d. shading.

It is important to note these modes are not discrete control inputs and are only meaningful in combination. For example, flapping involves both wing lifting and spreading (modes 1 and 2), but neither mode alone represents a full wingbeat. Decomposing these overlapping contributions into two modes, however, allows us to measure how they vary in timing and magnitude within and across wingbeats (Fig. 3 & 4), an insight difficult to discern from traditional kinematic parameters such as wingtip trajectory or angle of attack changes [27, 5].

## Morphing Signatures

While shared patterns across the hawks exist, the high variation should be noted (Fig. 3). Across our results we do not find stereotyped, repeatable shape changes. As shown in previous research, flight control is highly flexible and shaped by internal-state, external conditions, learned experience, and individual variation [28, 29, 19, 17, 30]. Although the hawks share core morphing modes, the individuals show distinct morphing signatures, differing in technique when facing different experimental conditions. Here we showcase examples split by individuals over repeated flights to further show the flexibility, adaptive variation in morphing flight (Fig. 4). There are subtle differences in flight maneuvers, between individuals (top left), and the same individual at different life stages (juvenile versus adult, bottom left). Experimental conditions further influenced morphing strategies. Introducing an obstacle midway between perches was accompanied by higher wing sweep scores, M-folding at the wrist, and tail shape changes (top right). Similarly, the added weight condition, a worn IMU, showed shorter glide phases and lower wing pitch scores (bottom right). These findings may also have implications for field studies using worn equipment, where even minor added mass measurably altered flight dynamics [31, 32].

**Figure 4.**
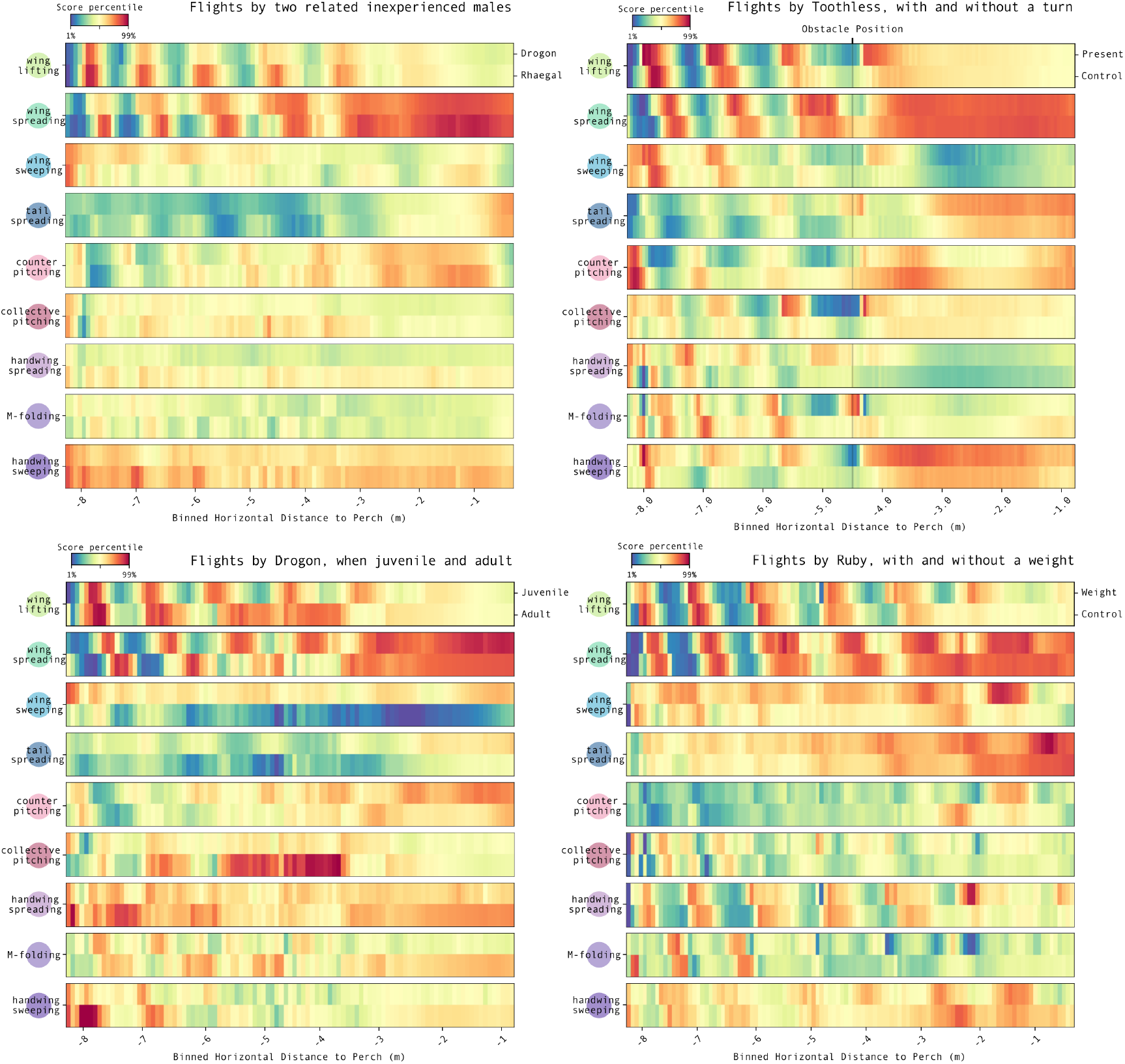
Morphing signatures across flight conditions. Individual hawks adapt flight in response to their environment and experience. Projecting into the kinematic morphospace makes direct comparison readily interpretable, even for subtle shape changes. Each heatmap shows multiple flights by the same hawk projected into a morphing shape mode (named on the left) and binned by distance to the perch (bin size = 0.05m). Colors show direction from the mean, for example wings lifting up (red) or down (blue) or held neutral (yellow). Experimental conditions or individuals are stacked for comparison (labeled on the top right). Top left: Related, inexperienced males (“Drogon”, top; “Rhaegal”, bottom) show distinct morphing signatures despite matched age, morphology, and experience (N=76 and 114 flights). Bottom left: The same hawk (“Drogon”) shows altered morphing with age: juvenile (<1yr, N=76) vs. adult (>3yr, N=46) under identical conditions. As an adult, his glide is longer, with more swept wings, and added tail raising mid-flight. Top right: In obstacle flights (male “Toothless”; N=30 with, 67 without), flap frequency and tail spreading is modified on approach to the obstacle and hand-wing adjustments increase with tail banking. Recovery is marked by increased wing sweep post-obstacle. Bottom right: Worn equipment (<4% body weight) alters flight in female “Ruby” (N=18 with, 48 without). Gliding decreases, wing pitch shifts, and changes in compensatory hand-wing morphing.

As well as coordination between the wings and tail, left-right couplings are also critical in flight control. We quantified the bilateral symmetry and asymmetry during morphing flight using the projection scores. We found major modes (wing lifting, spreading, and sweeping) were highly symmetrical during both flapping acceleration from take-off and in glides preceding landing. In contrast, other shape change modes showed greater flexibility and variation in asymmetry by flight context. During acceleration, fine-tune morphing was more symmetrical, likely aiding in stable, straight powered flight. The greater asymmetry recorded during gliding and deceleration is then likely important for maneuverability and landing precision [19].

Bird flight has evolved to overcome environmental and morphological challenges, where there is no single optimal flight configuration nor morphing strategy [23]. The true control space of neuromuscular activations is not defined here, but the kinematic morphospace provides a proxy for the resultant shape configurations without assumptions about joint angles or control inputs. In addition, aerodynamic function cannot be inferred from shape change alone, but these shape modes provide a robust foundation for testing hypotheses in bio-inspired robotics and control [33, 34, 35, 36]. Morphing shape modes offer a biologically meaningful basis for studying adaptive flight strategies, even without explicit mappings from neural input to motion. Decompositions into the temporal and spatial elements of morphing flight could also complement future analyses.

## Projection to Different Species

Morphing shape modes are linear combinations of 3D positions, making reconstructions directly interpretable and generalizable across morphologies and species. While transforming PCA loadings into a new morphology breaks the strict orthogonality of the original basis, the resulting shape modes remain biologically interpretable. These transformed axes no longer represent independent biomechanical degrees of freedom, but they nonetheless offer a consistent descriptive framework for comparing morphing strategies across species.

As a proof of concept, we projected Harris’ hawk flight into the morphologies of other birds using cadaver-based landmarks (Fig. 5). These projections are not intended to represent true flight postures in other species, but to demonstrate how shared shape modes can define a common coordinate space for cross-species analysis. Where traditional kinematics struggles to support meaningful comparison across different morphologies, morphing shape modes provide a shared descriptive basis. This framework distinguishes between common axes of variation and species- or individual-specific shape change strategies, enabling intuitive, high-sensitivity comparisons and suggesting new directions for bio-inspired design.

**Figure 5.**
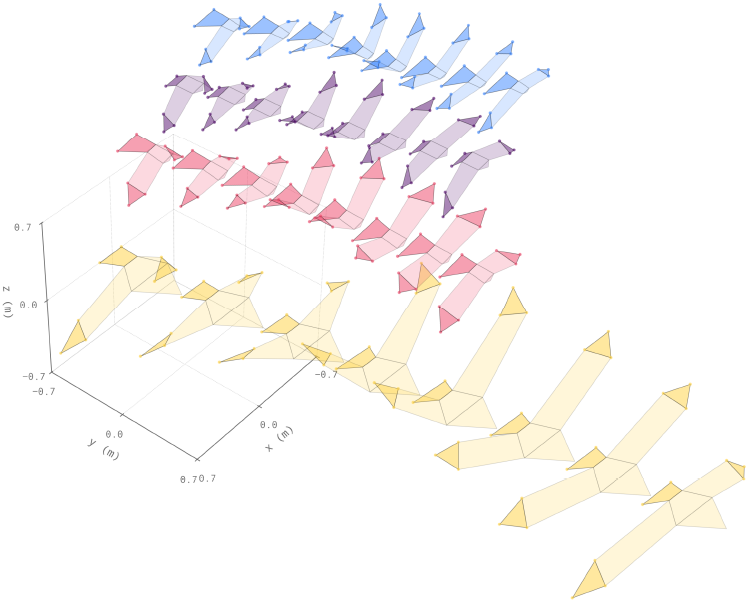
Cross-species application of kinematic morphospace. Morphing shape modes and scores derived from Harris’ hawks projected to three other species: Peregrine falcon (top, blue), Mallard (upper-mid, dark purple), and American pelican (bottom, yellow), alongside the original hawk (lower-mid, red). Each performs the same sequence of morphing shape changes (upstroke followed by downstroke) reconstructed within its own morphology. Transformations were based on homologous flight feather landmarks using cadaver measurements from Harvey et al. 2022. While PCA modes are orthogonal in the hawk kinematic morphospace, projecting onto new morphologies via transformed loadings breaks this orthogonality. Nevertheless, the resulting shape changes remain biologically interpretable. Future scores recorded from flight in other species should therefore be treated as descriptive positions within a shared space, rather than as independent biomechanical axes. Offsets added for clarity.

By reducing complex morphing into biologically meaningful components, we offer a generalizable framework for quantifying and comparing high-dimensional movement across behaviors, individuals, and species. Our approach captures coordination, symmetry, and adaptive response under varying conditions, distinguishes major and minor variation, and remains sensitive to subtle, behaviorally relevant differences. More broadly, it establishes a foundation for studying natural motion through shared shape spaces—structured, biologically grounded representations of movement with applications in biomechanics, evolutionary biology, robotics, animation, and bio-inspired engineering.

## Author Contributions

Conceptualization: LAF, GKT

Methodology: LAF, JS, MKH, CHB

Investigation: LAF, JS, MKH, CHB

Data Curation: LAF

Software: LAF

Formal Analysis: LAF

Visualization: LAF

Funding acquisition: GKT

Resources: GKT

Supervision: GKT

Writing – original draft: LAF

Writing – review & editing: GKT, JS, MKH, CHB

## Acknowledgments

This project has received funding from the European Research Council (ERC) under the European Union’s Horizon 2020 research and innovation programme (grant agreement no. 682501). The work of L.A.F. was supported by funding from the Biotechnology and Biological Sciences Research Council (BBSRC; grant no. BB/M011224/1) through the Interdisciplinary Bioscience Doctoral Training Partnership, and supported by Schmidt Sciences, LLC.,. We thank M. Parker, H. Sanders, and L. Larkman for animal husbandry and handling during experiments. We thank Kirsty Yeomans for her illustrations.

